# Cross-sectional association between blood cholesterol and calcium levels in genetically diverse strains of mice

**DOI:** 10.1101/2023.02.08.527123

**Authors:** Cody M. Cousineau, Kaelin Loftus, Gary A. Churchill, Dave Bridges

## Abstract

Genetically diverse outbred mice allow for the study of genetic variation in the context of high dietary and environmental control. Using a machine learning approach we investigated clinical and morphometric factors that associate with serum cholesterol levels in 840 genetically unique mice of both sexes, and on both a control chow and high fat high sucrose diet. We find expected elevations of cholesterol in male mice, those with elevated serum triglycerides and/or fed a high fat high sucrose diet. The third strongest predictor was serum calcium which correlated with serum cholesterol across both diets and sexes (r=0.39-0.48). This is in-line with several human cohort studies which show associations between calcium and cholesterol, and calcium as an independent predictor of cardiovascular events.

## Introduction

Elevated blood cholesterol, particularly in the forms of atherogenic LDL particles are causal of cardiovascular disease, the major cause of death in Western societies [1]. Cholesterol levels in humans vary widely depending on multiple factors including genetics, diet and other lifestyle factors, with genetics and lifestyle each contributing roughly equally to cardiovascular disease risk [2].

Human genetics has led to a sophisticated understanding of how cholesterol synthesis and excretion is regulated, and how this varies between individuals and diets [3]. Understanding complex genetic traits like cholesterol is a challenge in human studies where diet and lifestyle data is subject to substantial instrument error. Here genetically diverse panels of mice, where the diet and environment are tightly controlled, and genetics can be well defined can provide solutions. Among several resources, the Diversity Outbred [4] and UM-HET3 [5] mice represent genetically diverse outbred populations, while the Collaborative Cross [6] and BXD [7] resources provide data on recombinant inbred mice. Collectively these resources are valuable tools to understand complex traits such as cholesterol.

In this study we performed a secondary data analysis of Diversity Outbred mice to understand clinical and phenotypic correlates of elevated cholesterol levels. Using a machine learning approach, we demonstrate that sex, diet, and serum triglycerides are strong predictors of cholesterol levels in these mice, but also that even with high dietary control serum calcium provides and independent correlate of serum cholesterol levels.

## Methods

### Diversity Outbred Data

The phenotype data for diversity outbred mice (RRID:IMSR_JAX:009376) contains data on 840 mice from the diversity outbred collection of both sexes [8]. The dataset includes at data for 162 phenotypes, measured once, twice, or weekly in the case of body weights. At weaning mice were placed on a high fat high sucrose diet (HFHS; Harlan TD.08811), or kept on a normal chow diet (NCD; LabDiet 5K52). In the final dataset there were 225 female mice on NCD, 224 male mice on NCD, 198 female mice on HFHS, and 193 male mice on HFHS. Cholesterol, triglycerides, and calcium were quantified in plasma using the Beckman Synchron DXC600Pro Clinical chemistry analyzer. Body composition by dual x-ray absorbitrometry (DEXA) on Lunar PIXImus densitometer (GE Medical Systems). Additional details in [9]

### BXD Data

Calcium and cholesterol levels from male and female BXD were described in [10]. These data were downloaded from GeneNetwork (http://www.genenetwork.org/) [11,12] as datasets BXD_12844, BXD_12914, BXD_12951, and BXD_12881. These datasets included 17 female and strains (72 mice) and 36 male strains (254 mice; RRID:MGI:2164899). These mice were maintained on a chow diet (SAFE; D04) with blood collected at 14 weeks of age. Data were averaged and analyzed by strain and sex.

### Statistics

All statistical analyses were performed using R version 4.2.0 [13]. Cholesterol data were not normally distributed within groups (p<0.05 by sex and diet stratified Shapiro-Wilk tests), so non-parametric pairwise tests were used. Summarized data is reported as mean +/− standard error of the mean. For all comparisons sex was first tested as a modifier, and then as a covariate. If there was significant evidence of sex modification, pairwise sex-stratified analyses are also reported. Regression trees were generated using the rpart package (version 4.1.19; [14]), and pruned based on the number of branches at the minimum cross-validated standard error rate. Statistical significance was set at an alpha of 0.05. All data and reproducible code are available for this manuscript at https://github.com/BridgesLab/PrecisionNutrition.

## Results

### Diversity outbred mice exhibit diet and sex dependent variation in cholesterol levels

We first evaluated the cholesterol levels in the diversity outbred mice measured at 8 and 19 weeks. Cholesterol levels for each group were similar at both time points (p=0.474 by pairwise Wilcoxon test, see Supplementary Figure 1). This indicates that cholesterol levels are stable between both time points. We stratified cholesterol levels by sex and diet. Via multivariate regression, we found the expected cholesterol elevations in mice on a HFHS diet (33.7 +/− 2.0 mg/dL, p=1.4 x 10^-56^), and male sex (16.9 +/− 2.0 mg/dL, p=3.0 x 10^-17^; Figure 1A). There was no evidence of a significant interaction between diet and sex (p=0.636).

**Figure 1:**
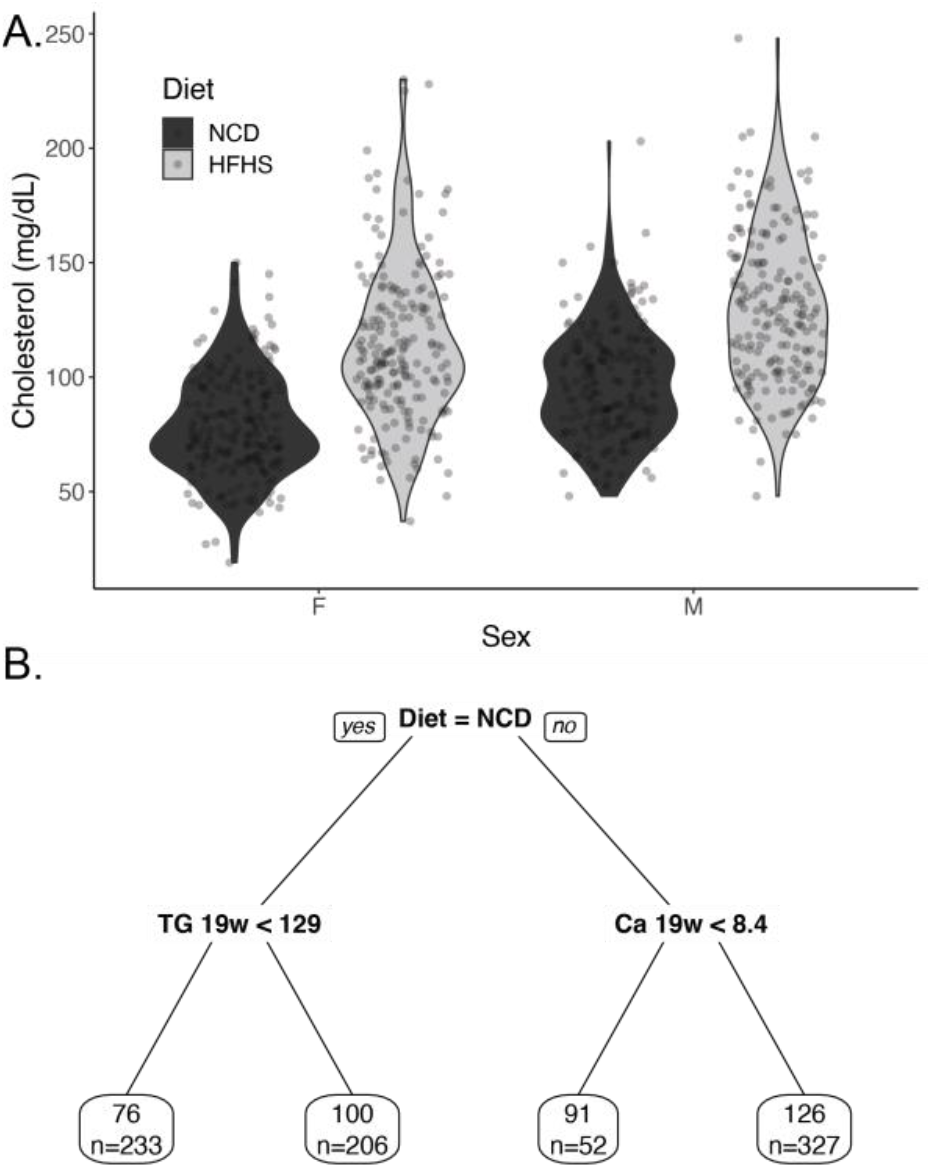
Description of cholesterol levels in diversity outbred mice. A) Violin plot of cholesterol levels of diversity outbred at 19 weeks, mice stratified by diet and sex. B) Pruned regression predicting cholesterol at 19 weeks. Above the box is the algorithmically generated cutoff for triglycerides (abbreviated TG in mg/dL), calcium (Ca in mg/dL), and body weight (BW in g). Each predictor was a phenotype measured at 19 weeks. Within each box, the value represents the average cholesterol level in that group (in mg/dL) and the number of mice in that group (n=822 in total).

### Diet, triglycerides and calcium associate with cholesterol levels

To define other potential associations between cholesterol and measured phenotypes in this dataset we generated a regression tree using the 165 phenotypes in this dataset (Figure 1B). The major classifier of cholesterol levels was the diet, and the second was triglycerides measured at 19 weeks. Serum calcium measured at 19 weeks was the third phenotype that associated with cholesterol levels, and body weight measured at 19 weeks was the fourth (Figure 1B).

Dyslipidemia often includes elevations of both triglyceride and cholesterol levels in both mice and humans, so the association of triglycerides with cholesterol was not unexpected. Via multivariate modelling accounting for the effects of diet and sex, a 100 mg/dL increase in triglycerides was associated with a 17.7 +/− 1.7 mg/dL elevation in cholesterol (p=3.8 x 10^-24^, Figure 2A).

**Figure 2:**
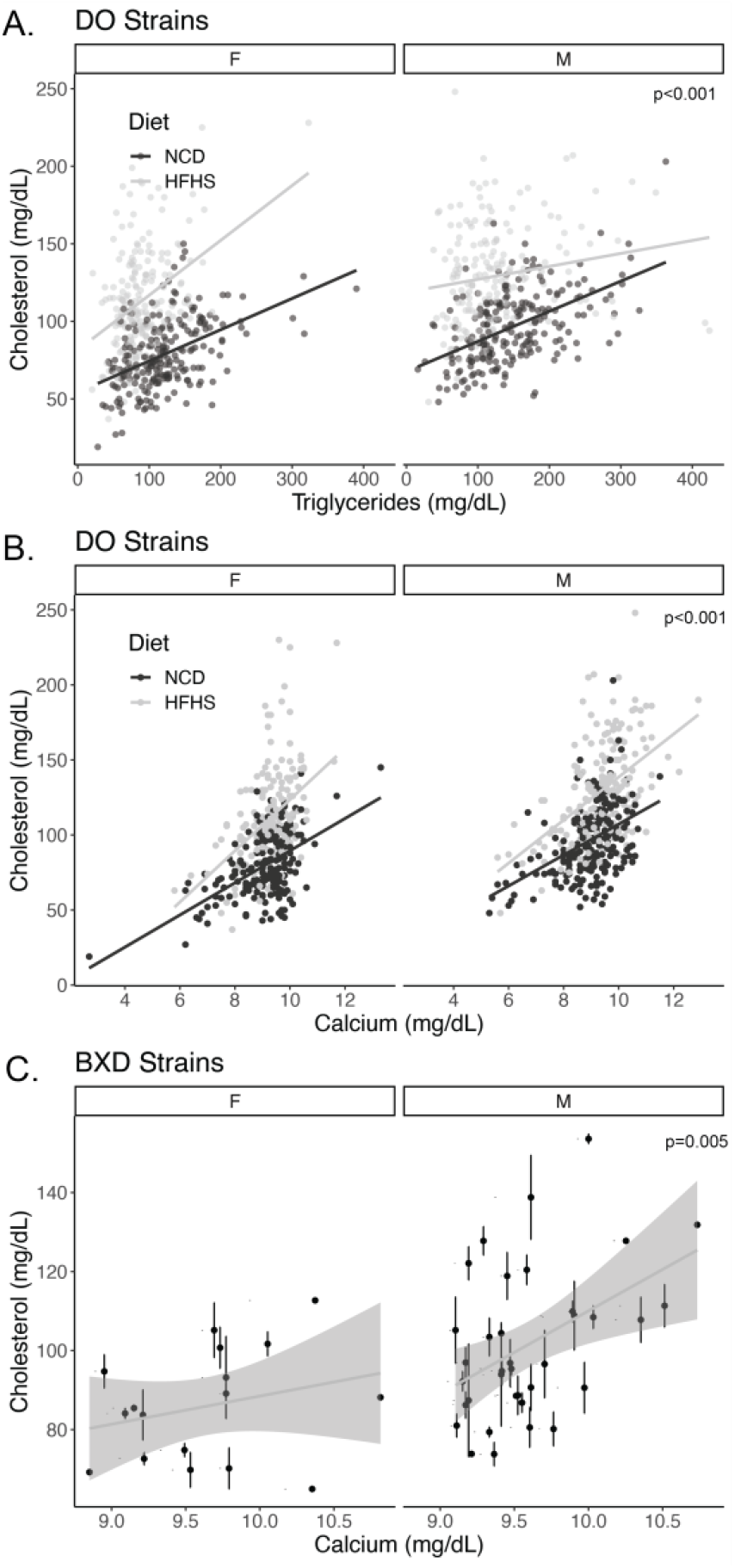
Cross-sectional associations of cholesterol with triglycerides and calcium. Sex and diet stratified scatter plots of A) triglyceride and B) calcium relationships with cholesterol levels at 19 weeks in diversity outbred mice (n=822 mice and strains). C) Cholesterol and calcium associations in male and female BXD strains (n=326 mice from 17 female and 36 male strains; error bars represent within-strain standard error of the mean). P-values indicate the level of significance for the diet and sex adjusted relationship between cholesterol and triglycerides or calcium from a multivariate linear model. The lines represent independent estimates for each group.

The strong cross-sectional association of calcium with cholesterol was not predicted by our research team. As shown in Figure 2B and Table 1, after adjusting for diet and sex, a one mg/dL increase in calcium is associated with a 12.7 +/− 0.8 mg/dL increase in cholesterol (p=3.0 x 10^-43^). Linear models predicting serum cholesterol including sex and diet, and calcium as covariates had an adjusted R^2^ of 0.45 with a partial effect size of 0.22 for serum calcium. We performed sub-group analyses and found that each diet-sex combination had broadly similar estimates for Spearman’s rho (ranging from 0.39 for HFHS females to 0.48 for HFHS males), each of which had a p-value of less than 2.2 x 10^-7^).

To externally test these findings, we evaluated a distinct dataset of genetically diverse mice, the BXD mouse collection. In a secondary data analysis using data in [10], we replicated this finding, showing a cross-sectional association between cholesterol and calcium levels after adjusting for sex differences. Similar to the data from the diversity outbred mice, we estimate a 14.8 +/− 5.1 mg/dL increase in cholesterol was observed for every 1 mg/dL increase in calcium (p=0.005). In the smaller BXD dataset there appeared to be a stronger relationship in male mice (Spearman’s rho=0.393 p=0.018, n=36 strains) than female mice (rho=0.269; p=0.297, n=17 strains), but in multivariate modelling there was no significant modifying effect of sex (p=0.178, likely due to the relatively small number of female strains).

In the diversity outbred mice, serum calcium levels are not significantly altered by sex (p=0.59), and only modestly increased by HFHS diets (0.30 +/− 0.07 mg/dL; p=5.1 x 10^-5^; Supplementary Figure 2A). Since calcium is normally tightly controlled by homeostatic mechanisms regulating calcium absorption and bone remodeling, we tested whether bone mineral content and density in these mice was associated with cholesterol levels. As shown in Supplementary Figure 2B and C there was no evidence of an association between bone mass or density measured at 21 weeks and cholesterol levels measured at 19 weeks (p=0.93 and 0.90 respectively) in the diversity outbred dataset.

## Discussion

In this study we report an association between calcium and cholesterol in two distinct mouse datasets. This relationship was similar across both sexes and over both normal chow and obesogenic high fat, high sucrose diets. The novel association between calcium and cholesterol study is strengthened by the expected findings that cholesterol is elevated in mice of male sex, with high triglycerides and fed HFHS diets. The finding that this association is largely independent of sex, diets or triglyceride levels suggests that serum calcium may represent a novel biomarker of dysfunctional cholesterol homeostasis and may point to novel mechanisms by which cholesterol could be controlled.

To our knowledge this is the first demonstration of an association between serum calcium and cholesterol in rodents. That being said, several large cross-sectional studies have consistently demonstrated a correlation between serum calcium and cholesterol in multiple populations [15–28]. In addition, calcium is also an longitudinal predictor of cardiovascular events in humans independent of BMI or blood pressure [17,28–34].

The present study does not speak to the directionality of this association, but there are some hints in the literature. A meta-analysis by demonstrates a 31% increased risk of myocardial infarction in patients with calcium supplementing with calcium compared to placebo [35]. Patients with primary hyperparathyroidism have elevated parathyroid hormone and calcium levels and are an interesting population to examine. The results of case-control studies evaluating cholesterol levels are mixed and there is limited evidence that PTH causes elevated cholesterol levels. While reports show that these patients also have significantly elevated total and/or LDL-cholesterol [36,37], though most others show either non-significant effect or even decreases [38–44].

In terms of whether cholesterol could be driving hypercalcemia, there is some positive evidence. Two interventional studies using statins have showed a reductions in calcium levels [45,46], whereas another single-arm study showed declines that did not reach significance [47]. A Mendelian Randomization approach using LDL-C as the instrument was also associated with elevated calcium levels [48], further supporting a potential causal relationship with cholesterol driving calcium levels. As this is a cross-sectional associative relationship we are not able at present to define directionality, much less the underlying biological mechanism(s). Whether calcium can modify serum cholesterol, or cholesterol can modify calcium are both important nutritional and pathophysiological questions, and future controlled mouse studies should shed light on the directions and mechanisms explaining this association.

There are several other strengths of this study. We present data on a large number of mice roughly equally divided between sexes and two diets and find consistent results across all groups. We have exceptional control of confounders such as diets, environment, activity levels, and other exposures that could affect the interpretation of the human studies. Our supervised machine learning approach used a large number of measured phenotypes to predict calcium levels, and set cutoffs in a data-driven manner. We consider the support of data from two independent genetically diverse mice populations another strength of this work. Calcium-cholesterol associations appear to be robust over a wide variation in genetics and not restricted to findings in inbred mouse populations. As such, this relationship holds over multiple diets, sexes, investigators, sites, and genetic backgrounds.

### Limitations of the present study

While there were multiple measurements of calcium and cholesterol in this dataset (at week 8 and week 19, after 5 and 16 weeks of HFHS/NCD respectively), cholesterol levels were stable through at these points. Therefore, it was possible to effectively evaluate the longitudinal association between cholesterol and calcium. In addition, this cross-sectional association does ascribe a directionality to this relationship, at this stage we think it plausible that calcium may increase cholesterol, that cholesterol might increase calcium, or that a third, unmeasured factor drives both factors. As this is a secondary data analysis, we are unable to evaluate differences in calcium-regulatory hormones, which we predict would vary more than the relatively homeostatic blood calcium levels. Another limitation is that cholesterol homeostasis is substantially different in mice and humans, especially in the fraction of cholesterol present in the HDL versus LDL fractions, due to the absence of CETP in mice. These data therefore largely reflect associations between calcium and the HDL pool.

In conclusion, in this work we use a machine learning approach to describe that diet, sex, triglyceride levels and calcium all contribute independently to the serum cholesterol levels in Diversity Outbred mice, and that the relationship between calcium and cholesterol holds true in BXD mice. These data support that the observed human relationships between serum and cholesterol levels are true in mice, and present an opportunity for further physiological and genetic dissection of this relationship.

## Acknowledgements

We would like to thank the members of the Bridges Laboratory for helpful discussions regarding this work. We would also like to acknowledge funding from the National Institutes of Diabetes and Digestive Kidney Diseases (NIDDK; R01DK107535 to DB), the National Institutes of General Medical Sciences (NIGMS R01GM07068) and the Undergraduate Research Opportunity Program (UROP to KL). We would also like to thank the developers and funders of the Diversity Outbred Database, Diversity Informatics Platform and GeneNetwork for their commitment to open science and for providing the data used in these analyses.

## Author Contributions

DB and CMC conceptualized this research study, decided and validated the methodologies, performed the investigations, wrote the original draft, the data, and prepared visualizations. Formal analyses were done by CMC, DB and KL. Data was provided by GAC. This work was administered and supervised by DB who along with GAC performed the data validation. Funding for this work was acquired by DB and GAC. All authors have read and agreed to the final published work.

## Notes

https://github.com/BridgesLab/PrecisionNutrition

### Competing Interest Statement

The authors have declared no competing interest.

## References

1. Grundy, S.M.; Stone, N.J.; Bailey, A.L.; Beam, C.; Birtcher, K.K.; Blumenthal, R.S.; Braun, L.T.; de Ferranti, S.; Faiella-Tommasino, J.; Forman, D.E.; et al. 2018 AHA/ACC/AACVPR/AAPA/ABC/ACPM/ADA/AGS/APhA/ASPC/NLA/PCNA Guideline on the Management of Blood Cholesterol: A Report of the American College of Cardiology/American Heart Association Task Force on Clinical Practice Guidelines. Circulation 2019, 139, e1082–e1143, doi:10.1161/CIR.0000000000000625.

2. Khera, A.V.; Emdin, C.A.; Drake, I.; Natarajan, P.; Bick, A.G.; Cook, N.R.; Chasman, D.I.; Baber, U.; Mehran, R.; Rader, D.J.; et al. Genetic Risk, Adherence to a Healthy Lifestyle, and Coronary Disease. New England Journal of Medicine 2016, 375, 2349–2358, doi:10.1056/NEJMoa1605086.

3. Khera, A.V.; Kathiresan, S. Genetics of Coronary Artery Disease: Discovery, Biology and Clinical Translation. Nature Reviews Genetics 2017, 18, 331–344, doi:10.1038/nrg.2016.160.

4. Churchill, G.A.; Gatti, D.M.; Munger, S.C.; Svenson, K.L. The Diversity Outbred Mouse Population. Mammalian Genome 2012, 23, 713–718, doi:10.1007/s00335-012-9414-2.

5. Jackson, A.U.; Fornés, A.; Galecki, A.; Miller, R.A.; Burke, D.T. Multiple-Trait Quantitative Trait Loci Analysis Using a Large Mouse Sibship. Genetics 1999, 151, 785–795, doi:10.1093/genetics/151.2.785.

6. Churchill, G.A.; Airey, D.C.; Allayee, H.; Angel, J.M.; Attie, A.D.; Beatty, J.; Beavis, W.D.; Belknap, J.K.; Bennett, B.; Berrettini, W.; et al. The Collaborative Cross, a Community Resource for the Genetic Analysis of Complex Traits. Nature genetics 2004, 36, 1133–1137, doi:10.1038/ng1104-1133.

7. Ashbrook, D.G.; Arends, D.; Prins, P.; Mulligan, M.K.; Roy, S.; Williams, E.G.; Lutz, C.M.; Valenzuela, A.; Bohl, C.J.; Ingels, J.F.; et al. A Platform for Experimental Precision Medicine: The Extended BXD Mouse Family. Cell Systems 2021, 12, 235–247.e9, doi:10.1016/j.cels.2020.12.002.

8. Gatti, D.M.; Simecek, P.; Somes, L.; Jeffery, C.T.; Vincent, M.J.; Choi, K.; Chen, X.; Churchill, G.A.; Svenson, K.L. The Effects of Sex and Diet on Physiology and Liver Gene Expression in Diversity Outbred Mice. bioRxiv 2017, 098657, doi:10.1101/098657.

9. Svenson, K.L.; Gatti, D.M.; Valdar, W.; Welsh, C.E.; Cheng, R.; Chesler, E.J.; Palmer, A.A.; McMillan, L.; Churchill, G.A. High-Resolution Genetic Mapping Using the Mouse Diversity Outbred Population. Genetics 2012, 190, 437–447, doi:10.1534/genetics.111.132597.

10. Andreux, P.A.; Williams, E.G.; Koutnikova, H.; Houtkooper, R.H.H.; Champy, M.-F.F.; Henry, H.; Schoonjans, K.; Williams, R.W.; Auwerx, J. Systems Genetics of Metabolism: The Use of the BXD Murine Reference Panel for Multiscalar Integration of Traits. Cell 2012, 150, 1287–1299, doi:10.1016/j.cell.2012.08.012.

11. Mulligan, M.K.; Mozhui, K.; Prins, P.; Williams, R.W. GeneNetwork: A Toolbox for Systems Genetics. In Methods in Molecular Biology; 2017; Vol. 331, pp. 75–120 ISBN 978-1-4939-6427-7.

12. Sloan, Z.; Arends, D.; Broman, K.W.; Centeno, A.; Furlotte, N.; Nijveen, H.; Yan, L.; Zhou, X.; Williams, R. W.; Prins, P. GeneNetwork: Framework for Web-Based Genetics. Journal of Open Source Software 2016, 1, 25, doi:10.21105/joss.00025.

13. R Core Team R: A Language and Environment for Statistical Computing 2019.

14. Therneau, Terry; Atkinson, Beth Rpart: Recursive Partitioning and Regression Trees.

15. Lind, L.; Jakobsson, S.; Lithell, H.; Wengle, B.; Ljunghall, S. Relation of Serum Calcium Concentration to Metabolic Risk Factors for Cardiovascular Disease. BMJ 1988, 297, 960–963.

16. Meng, X.; Han, T.; Jiang, W.; Dong, F.; Sun, H.; Wei, W.; Yan, Y. Temporal Relationship Between Changes in Serum Calcium and Hypercholesteremia and Its Impact on Future Brachial-Ankle Pulse Wave Velocity Levels. Frontiers in Nutrition 2021, 8.

17. Jorde, R.; Sundsfjord, J.; Fitzgerald, P.; Bønaa, K.H. Serum Calcium and Cardiovascular Risk Factors and Diseases. Hypertension 1999, 34, 484–490, doi:10.1161/01.HYP.34.3.484.

18. Kennedy, A.; Vasdev, S.; Randell, E.; Xie, Y.-G.; Green, K.; Zhang, H.; Sun, G. Clinical Medicine: Endocrinology and Diabetes: Abnormality of Serum Lipids Are Independently Associated with Increased Serum Calcium Level in the Adult Newfoundland Population. Clinical medicine. Endocrinology and diabetes 2009, 2, CMED.S2974, doi:10.4137/CMED.S2974.

19. Saltevo, J.; Niskanen, L.; Kautiainen, H.; Teittinen, J.; Oksa, H.; Korpi-Hyövälti, E.; Sundvall, J.; Männistö, S.; Peltonen, M.; Mäntyselkä, P.; et al. Serum Calcium Level Is Associated with Metabolic Syndrome in the General Population: FIN-D2D Study. European Journal of Endocrinology 2011, 165, 429–434, doi:10.1530/EJE-11-0066.

20. Gallo, L.; Faniello, M.C.; Canino, G.; Tripolino, C.; Gnasso, A.; Cuda, G.; Costanzo, F.S.; Irace, C. Serum Calcium Increase Correlates With Worsening of Lipid Profile. Medicine (Baltimore) 2016, 95, e2774, doi:10.1097/MD.0000000000002774.

21. De Bacquer, D.; De Henauw, S.; De Backer, G.; Kornitzer, M. Epidemiological Evidence for an Association between Serum Calcium and Serum Lipids. Atherosclerosis 1994, 108, 193–200, doi:10.1016/0021-9150(94)90114-7.

22. Wilson, P.W.; Garrison, R.J.; Abbott, R.D.; Castelli, W.P. Factors Associated with Lipoprotein Cholesterol Levels. The Framingham Study. Arteriosclerosis: An Official Journal of the American Heart Association, Inc. 1983, 3, 273–281, doi:10.1161/01.ATV.3.3.273.

23. Chou, C.-W.; Fang, W.-H.; Chen, Y.-Y.; Wang, C.-C.; Kao, T.-W.; Wu, C.-J.; Chen, W.-L. Association between Serum Calcium and Risk of Cardiometabolic Disease among Community-Dwelling Adults in Taiwan. Sci Rep 2020, 10, 3192, doi:10.1038/s41598-020-60209-w.

24. Sabanayagam, C.; Shankar, A. Serum Calcium Levels and Hypertension Among US Adults. The Journal of Clinical Hypertension 2011, 13, 716–721, doi:10.1111/j.1751-7176.2011.00503.x.

25. Rohrmann, S.; Garmo, H.; Malmström, H.; Hammar, N.; Jungner, I.; Walldius, G.; Van Hemelrijck, M. Association between Serum Calcium Concentration and Risk of Incident and Fatal Cardiovascular Disease in the Prospective AMORIS Study. Atherosclerosis 2016, 251, 85–93, doi:10.1016/j.atherosclerosis.2016.06.004.

26. Green, M.A.; Jucha, E. Interrelationships between Blood Pressure, Serum Calcium and Other Biochemical Variables. International Journal of Epidemiology 1987, 16, 532–536, doi:10.1093/ije/16.4.532.

27. He, L.; Qian, Y.; Ren, X.; Jin, Y.; Chang, W.; Li, J.; Chen, Y.; Song, X.; Tang, H.; Ding, L.; et al. Total Serum Calcium Level May Have Adverse Effects on Serum Cholesterol and Triglycerides Among Female University Faculty and Staffs. Biol Trace Elem Res 2014, 157, 191–194, doi:10.1007/s12011-014-9895-9.

28. Lind, L.; Skarfors, E.; Berglund, L.; Lithell, H.; Ljunghall, S. Serum Calcium: A New, Independent, Prospective Risk Factor for Myocardial Infarction in Middle-Aged Men Followed for 18 Years. Journal of Clinical Epidemiology 1997, 50, 967–973, doi:10.1016/S0895-4356(97)00104-2.

29. Reid, I.R.; Gamble, G.D.; Bolland, M.J. Circulating Calcium Concentrations, Vascular Disease and Mortality: A Systematic Review. Journal of Internal Medicine 2016, 279, 524–540, doi:10.1111/joim.12464.

30. Foley, R.N.; Collins, A.J.; Ishani, A.; Kalra, P.A. Calcium-Phosphate Levels and Cardiovascular Disease in Community-Dwelling Adults: The Atherosclerosis Risk in Communities (ARIC) Study. American Heart Journal 2008, 156, 556–563, doi:10.1016/j.ahj.2008.05.016.

31. Slinin, Y.; Blackwell, T.; Ishani, A.; Cummings, S.R.; Ensrud, K.E.; MORE Investigators Serum Calcium, Phosphorus and Cardiovascular Events in Post-Menopausal Women. Int J Cardiol 2011, 149, 335–340, doi:10.1016/j.ijcard.2010.02.013.

32. Jorde, R.; Schirmer, H.; Njølstad, I.; Løchen, M.-L.; Bøgeberg Mathiesen, E.; Kamycheva, E.; Figenschau, Y.; Grimnes, G. Serum Calcium and the Calcium-Sensing Receptor Polymorphism Rs17251221 in Relation to Coronary Heart Disease, Type 2 Diabetes, Cancer and Mortality: The Tromsø Study. Eur J Epidemiol 2013, 28, 569–578, doi:10.1007/s10654-013-9822-y.

33. Walsh, J.P.; Divitini, M.L.; Knuiman, M.W. Plasma Calcium as a Predictor of Cardiovascular Disease in a Community-Based Cohort. Clin Endocrinol (Oxf) 2013, 78, 852–857, doi:10.1111/cen.12081.

34. Wang, M.; Yan, S.; Peng, Y.; Shi, Y.; Tsauo, J.-Y.; Chen, M. Serum Calcium Levels Correlates with Coronary Artery Disease Outcomes. Open Medicine 2020, 15, 1128–1136, doi:10.1515/med-2020-0154.

35. Effect of Calcium Supplements on Risk of Myocardial Infarction and Cardiovascular Events: Meta-Analysis | The BMJ Available online: https://www.bmj.com/content/341/bmj.c3691 (accessed on 4 February 2023).

36. Procopio, M.; Barale, M.; Bertaina, S.; Sigrist, S.; Mazzetti, R.; Loiacono, M.; Mengozzi, G.; Ghigo, E.; Maccario, M. Cardiovascular Risk and Metabolic Syndrome in Primary Hyperparathyroidism and Their Correlation to Different Clinical Forms. Endocrine 2014, 47, 581–589, doi:10.1007/s12020-013-0091-z.

37. Luigi, P.; Chiara, F.M.; Laura, Z.; Cristiano, M.; Giuseppina, C.; Luciano, C.; Giuseppe, P.; Sabrina, C.; Susanna, S.; Antonio, C.; et al. Arterial Hypertension, Metabolic Syndrome and Subclinical Cardiovascular Organ Damage in Patients with Asymptomatic Primary Hyperparathyroidism before and after Parathyroidectomy: Preliminary Results. Int J Endocrinol 2012, 2012, 408295, doi:10.1155/2012/408295.

38. Luboshitzky, R.; Chertok-Schaham, Y.; Lavi, I.; Ishay, A. Cardiovascular Risk Factors in Primary Hyperparathyroidism. J Endocrinol Invest 2009, 32, 317–321, doi:10.1007/BF03345719.

39. Ring, M.; Farahnak, P.; Gustavsson, T.; Nilsson, I.-L.; Eriksson, M.J.; Caidahl, K. Arterial Structure and Function in Mild Primary Hyperparathyroidism Is Not Directly Related to Parathyroid Hormone, Calcium, or Vitamin D. PLoS One 2012, 7, e39519, doi:10.1371/journal.pone.0039519.

40. Farahnak, P.; Lärfars, G.; Sten-Linder, M.; Nilsson, I.-L. Mild Primary Hyperparathyroidism: Vitamin D Deficiency and Cardiovascular Risk Markers. J Clin Endocrinol Metab 2011, 96, 2112–2118, doi:10.1210/jc.2011-0238.

41. Christensson, T.; Einarsson, K. Serum Lipids before and after Parathyroidectomy in Patients with Primary Hyperparathyroidism. Clinica Chimica Acta 1977, 78, 411–415, doi:10.1016/0009-8981(77)90074-2.

42. Ejlsmark-Svensson, H.; Rolighed, L.; Rejnmark, L. Effect of Parathyroidectomy on Cardiovascular Risk Factors in Primary Hyperparathyroidism: A Randomized Clinical Trial. The Journal of Clinical Endocrinology & Metabolism 2019, 104, 3223–3232, doi:10.1210/jc.2018-02456.

43. Hagström, E.; Lundgren, E.; Rastad, J.; Hellman, P. Metabolic Abnormalities in Patients with Normocalcemic Hyperparathyroidism Detected at a Population-Based Screening. European Journal of Endocrinology 2006, 155, 33–39, doi:10.1530/eje.1.02173.

44. Kaji, H.; Hisa, I.; Inoue, Y.; Sugimoto, T. Low Density Lipoprotein-Cholesterol Levels Affect Vertebral Fracture Risk in Female Patients with Primary Hyperparathyroidism. Exp Clin Endocrinol Diabetes 2010, 118, 371–376, doi:10.1055/s-0029-1224152.

45. Soh, J.F.; Bodenstein, K.; Yu, O.H.Y.; Linnaranta, O.; Renaud, S.; Mahdanian, A.; Su, C.-L.; Mucsi, I.; Mulsant, B.; Herrmann, N.; et al. Atorvastatin Lowers Serum Calcium Levels in Lithium-Users: Results from a Randomized Controlled Trial. BMC Endocr Disord 2022, 22, 238, doi:10.1186/s12902-022-01145-w.

46. Farhan, H.A.; Khazaal, F.A.; Mahmoud, I.J.; Haji, G.F.; Alrubaie, A.; Abdulraheem, Y.; * A.M.A.; Alkuraishi, M. Efficacy of Atorvastatin in Treatment of Iraqi Obese Patients with Hypercholesterolemia. AL-Kindy College Medical Journal 2014, 10, 62–69.

47. Montagnani, A.; Gonnelli, S.; Cepollaro, C.; Pacini, S.; Campagna, M.S.; Franci, M.B.; Lucani, B.; Gennari, C. Effect of Simvastatin Treatment on Bone Mineral Density and Bone Turnover in Hypercholesterolemic Postmenopausal Women: A 1-Year Longitudinal Study. Bone 2003, 32, 427–433, doi:10.1016/S8756-3282(03)00034-6.

48. Li, S.; Schooling, C.M. A Phenome-Wide Association Study of Genetically Mimicked Statins. BMC Med 2021, 19, 1–11, doi:10.1186/s12916-021-02013-5.

