## Supplementary Figure Legends for "Cross-sectional association between blood cholesterol and calcium levels in genetically diverse strains of mice"

**Supplementary Figure 1: Cholesterol levels are stable across time in diversity outbred mice.** Average cholesterol levels, and levels measured at 8 and 19 weeks, stratified by sex and diet.

**Supplementary Figure 2: Calcium is not strongly associated with diet, sex or bone mass/density in diversity outbred mice.** A) Violin plot of calcium levels at 19 weeks across diets and sex. Sex and diet stratified scatter plots of the relationships between bone mineral content (A) and bone density (B) via DEXA scan and their relationships with cholesterol levels at 19 weeks. For A, the p values represent the significance of diet and sex from a multivariate linear model. For B and C, p values indicate the significance for the diet and sex adjusted relationship between cholesterol and bone mineral content or density from a multivariate linear model.
