## Supplementary figures and images for "Cross-sectional association between blood cholesterol and calcium levels in genetically diverse strains of mice"

### Supplementary Figures

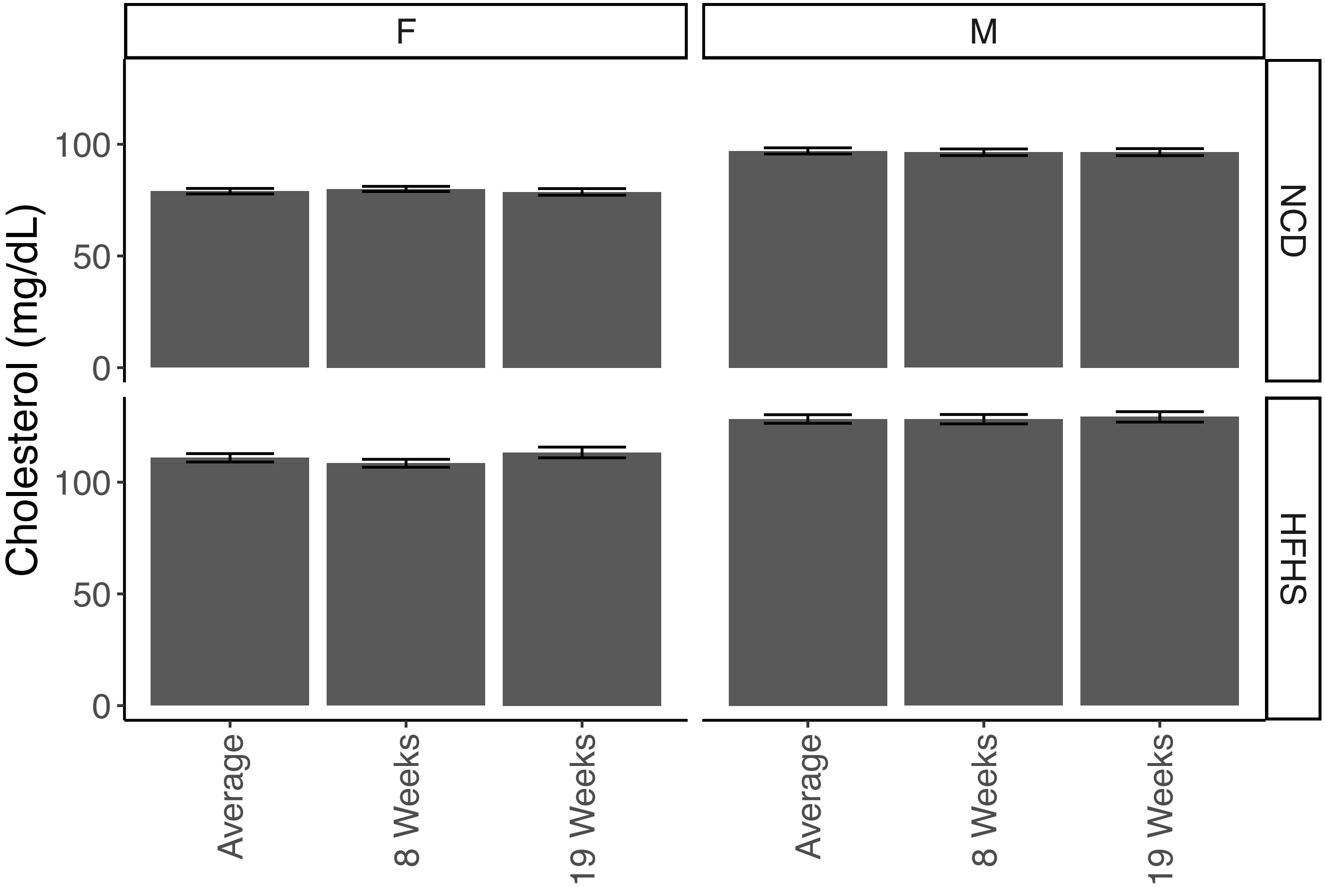

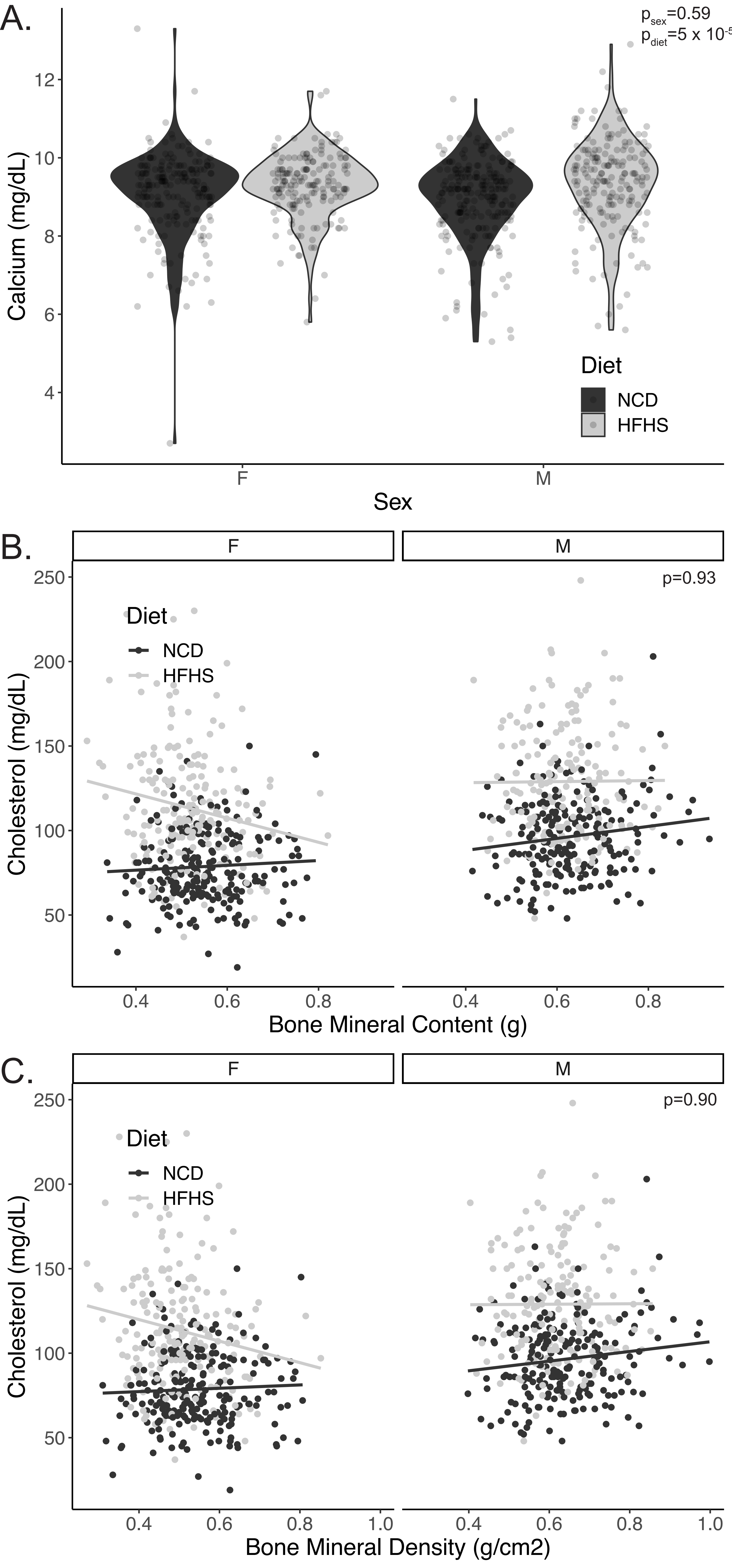
